# Computational search for UV radiation resistance strategies in *Deinococcus swuensis* isolated from Paramo ecosystems

**DOI:** 10.1101/734129

**Authors:** Jorge Díaz-Riaño, Leonardo Posada, Iván Acosta, Carlos Ruíz-Pérez, Catalina García-Castillo, Alejandro Reyes, María Mercedes Zambrano

## Abstract

Ultraviolet radiation (UVR) is widely known as deleterious for many organisms since it can cause damage to biomolecules either directly or indirectly via the formation of reactive oxygen species. The goal of this study was to analyze the capacity of high-mountain *Espeletia hartwegiana* plant phyllosphere microorganisms to survive UVR and to identify genes related to resistance strategies. A strain of *Deinococcus swuensis* showed a high survival rate of up to 60% after UVR treatment at 800*J/m*^2^ and was used for differential expression analysis using RNA-seq after exposing cells to 400*J/m*^2^ of UVR (with *>*95% survival rate). Differentially expressed genes were identified using the R-Bioconductor package NOISeq and compared with other reported resistance strategies reported for this genus. Genes identified as being overexpressed included transcriptional regulators and genes involved in protection against damage by UVR. Non-coding (nc)RNAs were also differentially expressed, some of which have not been previously implicated. This study characterized the early resistance strategy used by *D. swuensis* and indicates the involvement of ncRNAs in the adaptation to extreme environmental conditions.

## Introduction

Diverse natural and artificial environments exposed to extreme temperature, pressure and/or radiation conditions are attractive sources of microorganisms with exceptional phenotypic and genotypic properties. The high-mountain Paramo biome, similar to the tundra biome of high latitudes, consists of high-elevation areas subject to harsh environmental conditions. The Paramo biome has a high solar incidence that can induce damage by ultraviolet radiation (UVR) that represents a survival challenge for organisms [50]. Ionizing radiation and UVR affect organisms by damaging cellular components such as nucleic acids, proteins, and lipids [30]. The deleterious effect on cells is caused by direct damage to DNA, such as chromosomal lesions that introduce both double-strand breaks (DSBs) and single-strand breaks (SSBs), and damage due to pyrimidine dimerization and photoproducts that inhibit DNA replication and transcription [2]. Most of the damage, however, is caused indirectly by the production of reactive oxygen species (ROS), such as the chemically reactive superoxide and hydroxyl radicals that in turn affect various cellular constituents, including proteins [30].

The electromagnetic spectrum of UVR is divided into ultraviolet A (UVA) with wavelengths from 315-400 nm (6.31e-19 – 4.97e-19 *J/m*^2^s), ultraviolet B (UVB, from 280-315 nm, 7.10e-19 – 6.31e-19 *J/m*^2^s) and ultraviolet C (UVC, from 100-280 nm, 1.99e-18 – 7.10e-19 *J/m*^2^s) [64]. The UV electromagnetic spectrum covering UVB and UVC is considered ionizing radiation [51]. While UVC is absorbed by the ozone layer and the atmosphere, about 8% of UVA and 1% of UVB reach the Earth’s surface [51]. The harmful effects of UVR on cellular components depend on the wavelength: UVA can travel farther into tissues and contributes to ROS (damage to lipids, proteins, and DNA) whereas UVB produces direct breaks in the DNA structure (pyrimidine dimers) [62]. Even though UVC radiation is not present on the Earth’s surface, its bactericidal potential is used for studying UV sensitivity in bacteria with a high tolerance to UVB or UVA radiation [27].

Organisms resistant to radiation have been identified in all three domains of life. The mechanisms proposed to be involved in resistance to UVR vary and include strategies for DNA repair, protection against ROS using either enzymes or non-enzymatic antioxidative defenses, such as intracellular manganese and pigment production, protein folding and degradation systems [18]. Bacteria, with their diverse metabolic capacity, have an uncanny ability to survive under extreme conditions and colonize habitats that are inhospitable to other groups of organisms [11]. Different levels of resistance to UVR have been reported in diverse bacterial species, highlighting a wide variation in the response and a need for understanding the physiological, biochemical and mechanical responses that confer UV tolerance in bacteria [19]. Perhaps the most representative members of the extremely radiation-resistant bacteria belong to the family Deinococcaceae, which can survive exposure to ionizing radiation over 12,000 Gy (J/Kg), UVR over 1000*J/m*^2^ and can grow under harsh chronic irradiation of 50 Gy per hour [12]. *Deinococcus swuensis*, whose genome was recently published from a strain isolated from soil in South Korea, is also reported to have high resistance to UVR [32] [26].

Transcriptomic studies of *D. radiodurans* under radiation stress have shown induction of genes involved in DNA repair, cell recovery and antioxidative defenses [37]. An RNA-Seq analysis of *D. gobiensis* also showed induction of genes for DNA repair and regulation in response to UVR [63]. These studies, together with more recent work [57], also indicate differential expression of a subset of small and noncoding RNAs (sRNAs/ncRNAs), molecules that do not encode functional proteins but can play important roles in regulation of transcription and translation [61].

The differential expression of ncRNAs, upon UVR treatment suggests that these molecules could be important in triggering protective mechanisms, even though their precise role during the stress response to high doses of radiation still remains to be determined. A new hypothesis suggests that sRNAs could contribute to cellular post-exposure recovery because they would remain largely undamaged due to their small size [57]. Experimental evidence places these sRNAs into different metabolic pathways, such as response to changes in temperature, pH and other lethal stressors [59]. Recently reported sRNAs identified to be involved in radiation resistance are Y-RNAs, molecules that adopt specific secondary structures and bind to proteins known as Ro that are conserved in several organisms [29]. In *D. radiodurans* Y-RNAs were found to bind the Ro orthologue Rsr to form a ribonucleoprotein (Ro-RNP) complex that functions as an effective machinery for bacterial RNA degradation [8]. *D. radiodurans* was found to upregulate and accumulate Ro-RNPs in response to UVR and cells lacking the Ro protein had decreased survival following UV exposure [9].

In this study we hypothesized that microorganisms capable of resisting UVR should be present in locations exposed to high solar incidence, such as the Andean mountain high-altitude Paramo biome. Previous results indicated that the phyllosphere microbiota associated with *Espeletia sp.*, a plant endemic to the Paramo, contained diverse microbial communities and genes involved in resistance to UV and other stress conditions [50], and could thus provide insight into microbial resistance strategies. The main goal of this work was to isolate UV resistant microorganisms from this plant phyllosphere and study their resistance mechanisms through gene expression analysis. One bacterial strain identified as *D. swuensis* showed high resistance to UV exposure in laboratory settings and differential regulation of genes and sRNAs that provide clues to the early adaptation of *D. swuensis* to extreme environmental conditions, such as those found in high Andean ecosystems.

## Materials and methods

### Isolation of bacterial strains, culture conditions and characterization

Microorganisms were isolated from *Espeletia hartwegiana* leaf surfaces by first dislodging microbes, as previously reported [5], and then plating serial dilutions on R2A Agar (BD Difco, Franklin Lakes, NJ) and Tryptone soy agar (TSA, Oxoid), supplemented with 50 mg/ml Nystatin (Sigma-Aldrich, St. Louis, MO) to avoid fungal growth, when necessary. Plates were incubated at 25°C for 15 days and checked daily for growth. Colonies with distinct morphologies were re-streaked in the same growth media until pure colonies were obtained. Strains were characterized microscopically using Gram staining and taxonomic identification was done by analysis of the 16S rRNA gene or the ITS region for fungi. DNA was obtained by resuspending colonies in 1ml Tris 10mM (pH 8.0), adding 25*µ*l proteinase K (10mg/ml) and incubating at 55°C for 25 min. DNA was purified from 500*µ*l of this cell suspension using the MO BIO Microbial Ultraclean DNA Purification Kit (Qiagen, Germany). PCR amplifications were done using primers 27F (5’ AGAGTTTGATCMTGGCTCAG 3’) and 1492R (5’ TACGGYTACCTTGTTACGACTT 3’) for bacteria, in a 50*µ*l reaction volume containing 1*µ*l DNA template, 0.2*µ*M of each primer, 0.2 mM dNTPs, 2.5 mM MgCl_2_, 1X Buffer and 1.25 U of Taq DNA polymerase (CorpoGen, Colombia) and the following amplification conditions: 4 min at 94°C, 35 cycles of 30 s at 94°C, 45 s at 55°C, 1 min a 72°C, and a final extension of 10 min at 72°C. Primers ITS5 (5’ GGAAGTAAAAGTCGTAACAAGG 3’) and ITS4 (5’ TCCTCCGCTTATTGATATGC 3’) were used to amplify fungi as described above but using 0.3 *µ*M primers and PCR reactions of 2 min at 94°C, followed by 35 cycles of 60 s at 94°C, 60 s at 55°C, 1 min a 72°C, and a final extension of 5 min at 72°C. Sequencing was performed in an Applied Biosystems 3500 Genetic Analyzer. Forward and reverse reads were assembled and analyzed using Geneious 8.2, removing low quality nuclotides, and queried against the NCBI nucleotide database using BLAST.

### Screen for UV resistance and *D. swuensis* survival curve

Strains were grown overnight in 3ml Tryptone soy broth (TSB, Oxoid), washed 3 times with PBS, and 20*µ*l of nine 1:10 serial dilutions were spotted, in triplicate, on TSA medium, allowed to dry, and exposed to UVC in a UV hood to obtain a fluence rate from 50 to 800*J/m*^2^, as previously described [45] and determined using a radiometer with an LP 471 UVC probe (Delta Ohm, HD2302.0). Survival was determined by plating irradiated cultures on TSA medium to determine CFU/ml. Survival of *D. swuensis* was measured at various points along the growth curve using three replicate cultures that were first grown overnight and then diluted 1:100 in 100 ml fresh TSB medium, and incubated at 30°C, with continuous agitation at 150rpm. Samples were taken at 15, 24, 40, 48, and 72 hours and exposed to 800, 1600 and 2400*J/m*^2^ to determine survival (CFU/ml), as mentioned above.

### RNA extraction and sequencing

Triplicate 48-hour *D. swuensis* cultures were grown in 3ml TSB, diluted 1:100 into 100 ml fresh medium and re-grown for 24h. Ten ml of each 24h culture were transferred to a sterile Petri dish and submitted to 400*J/m*^2^ irradiation. After exposure, bacterial cells were immediately placed on ice, and centrifuged at 4600 x g for 15 minutes (4°C). Control aliquots from the same culture were not submitted to irradiation. After centrifugation, pellets were re-suspended in 1ml TriZol (Promega), lysed with Matrix B lysing beads (MP Biomedicals) in a FastPrep (MP Biomedicals) using 6.5 m/s for 40 seconds, and then centrifuged at 15,000 x g for 1 minute at 4°C. RNA in supernatants was recovered with the DirectZol RNA extraction kit (Zymo Research). Only RNA with a RIN *>*8 was used for sequencing at Macrogen (Seoul, Korea) on an illumina Hi-seq 2000, with 100 nucleotide paired-end reads.

### Preprocessing and mapping sequencing data

Quality control was made with FastQC (v.0.11.2) (http://www.bioinformatics.bbsrc.ac.uk/projects/fastqc/), Illumina adapters were trimmed with Trimmomatic (v.0.36) [6], rRNA depletion was performed with SortmeRNA (v.2.1) [28] using the Silva16S, 23S and 5S rRNA gene databases (release 128 downloaded on January 2017 from https://www.arb-silva.de/nocache/download/archive/release_128/) [47]. Sequences were mapped against the *D. swuensis* NCBI reference genome DY59 (accession number: GCF 000800395.1) [32] with Subread (V1.5.0-p3) [33], parameters included an insert size of 250 bp, a maximum of 3 mismatches and 5 indels. Features present in the reference annotation were extracted from the gff file and relative abundances were calculated with featureCounts, a tool included in the R package subread (v.1.5.0-p3) [34], and an in-house script. Remaining (unmapped) sequences were randomly subsampled (10%) and searched against the nt (nucleotide collection) database with Blastn and processed with MEGAN (v.6.9.4) at a threshold of 1*X*10^−5^ [24].

### Differential expression analysis

Annotated features from the reference genome such as coding DNA sequences (CDS’s), ncRNAs, pseudogenes, rRNAs and tRNAs were selected for analysis. The R-Bioconductor package NoiseqBio (2.18.0) [55] was used to measure differential gene expression between irradiated and non-irradiated control conditions. The workflow included a variance diagnostic (Cochran C test), analysis of sequencing depth, search for biases due to i) feature length and RNA amount and ii) detection of features with low counts, and the nonparametric analysis of differentially expressed features (based on Bayesian statistics). Counts were normalized to reads per kilobase of feature length per million mapped reads (RPKM) and by trimmed mean of M-values (TMM). Filtering of features with low counts was applied in order to remove those features that had an average expression of less than 5 CPM (counts per million) per condition and a variation coefficient higher than 100 in all conditions, which introduces noise and can lead to unreliable results for differential expression analysis [52]. Genes with a cut-off probability of expression above 0.8 and a log_2_ fold-change greater than or equal to 1.0 were considered as differentially expressed genes. Sequences coding for annotated hypothetical proteins were queried against the NCBI nr database using BLASTp and used for domain search with HMMER (V.3.1) [40] against the PFAM database [17].

### ncRNA computational analysis

Intergenic regions of the reference genome showing a significant number of mapped transcriptomic reads (minimum 6X coverage) were retrieved as potentially containing ncRNAs. Filters for the regions selected were based on the number of hits (read counts) and the region length (*>*50pb). Candidate regions were compared against the Rfam and NCBI nucleotide-nr databases [25] [48] using covariance models implemented in Infernal (V.1.1) [41]. All ncRNA candidates were processed for differential expression analysis using the workflow described above.

### Quantitative Real time PCR (qRT PCR) validation

Primers were designed using the IDT primerQuest tool (https://www.idtdna.com/Primerquest/Home/Index) to have a TM=*∼*60°C, a final amplified product size of *∼* 200pb and GC content *∼*50 % Table S1. RNA samples were quantified using a Qubit fluorometric system (Invitrogen) and used at the same concentrations for cDNA synthesis using Super script III reverse transcriptase (Invitrogen). qPCRs were run in a LightCycler ® 96 System (Roche) using the FastStart Essential DNA Green Master kit (Roche) and the following conditions: 1 cycle of 600 s at 95°C, then 45 cycles of 10 s at 95°C, 10s at the annealing temperature and a final extension at 72°C for 10s; a melting curve after the amplification confirmed a single peak and indicated a specific qPCR product. Relative expression was obtained by normalizing with the single copy gene QR90 RS09970 that codes for a succinate dehydrogenase, that showed similar expression levels among the different samples and conditions in the RNA-seq analysis, and the equation proposed by [46]. Primer efficiencies were determined using 1:10 serial dilutions of genomic *D. swuensis* DNA and the same PCR program described above.

## Results

### Strain isolation and radiation resistance

Microorganisms were isolated from the phyllosphere of *Espeletia* plants located in the National Park Los Nevados in Colombia [50]. Isolates with distinct colony morphologies were obtained by plating dilutions of the material dislodged from leaf surfaces on various media. Taxonomic identification using both 16S rRNA gene and ITS sequence analyses showed that this collection of isolates consisted of 10 fungi, 11 Gram-positive and 29 Gram-negative bacteria. To determine if any of these strains were resistant to UV radiation, as predicted for organisms living at these high-altitude ecosystems [50], all isolates were subjected to irradiation with UVC. A screen using varying levels of exposure, up to 800*J/m*^2^, showed that very few strains were capable of surviving these conditions. The most resistant strain was a bacterium identified as *D. swuensis* (strain CG1225), followed by the fungi *Cryptococcus flavescens* and *Rhodotorula mucilaginosa* Fig 1A. Other isolates showed reduced levels of resistance. Given that *D. swuensis* CG1225 showed the highest resistance to UVC exposure, with *>*60% survival at the highest dose tested (800*J/m*^2^), this strain was selected to further study its response to irradiation using RNAseq analysis.

**Fig 1.**
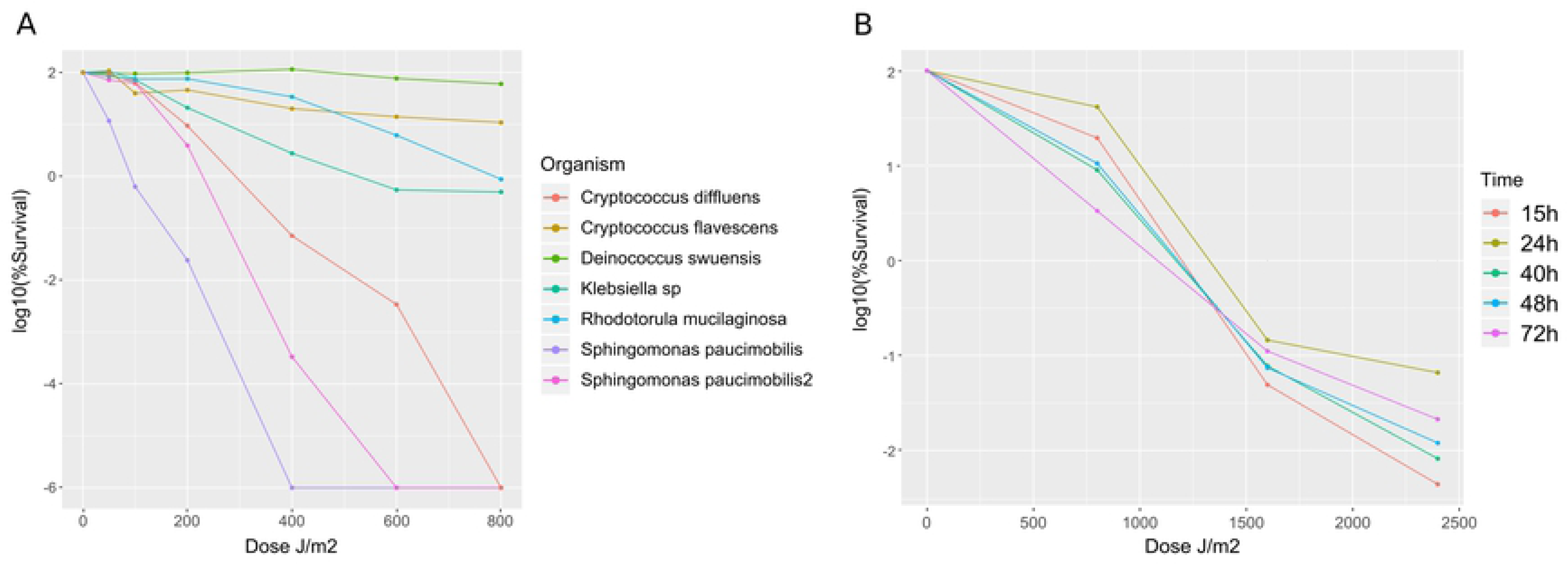
Survival to UV-C exposure of *Espeletia* phyllosphere-associated microorganisms. **(A)** Bacteria and yeast isolated from the plant phyllosphere were exposed to different UVR doses. Survival was measured as the percentage of CFUs obtained when compared to unexposed cells of the same strain. **(B)** Survival (mean *±* SD, n=3) of *D. Swuensis* harvested at different time points along the growth curve (15-72h) and exposed to different doses of UV-C.

In order to determine the best conditions for RNA extraction, a survival curve was first performed by harvesting *D. swuensis* cells at five different times along the growth curve and exposing these cells to varying doses of UVC, including doses above 800*J/m*^2^ used previously Fig 1B. Radiation resistance was similar for all time points examined along the growth curve (15 h to 72 h cultures), even up to the maximum exposure tested (2400*J/m*^2^). However, degradation of the extracted RNA was observed at increased doses of UV exposure. Thus we selected the treatment of 400*J/m*^2^ with 24 h cultures for subsequent RNA extractions to ensure sufficient recovery of high-quality RNA.

### Pre-processing and mapping sequencing data

Total RNA was obtained for three independent replicates of unexposed controls (C1, C2, and C3) and irradiated cultures (IR1, IR2, and IR3). RNA-seq was carried out using 100-nucleotide paired-end sequencing on an Illumina HiSeq. On average, 16.4 million reads were obtained per sample Table 1. After quality processing and adapter removal, rRNA filtering was performed using the SILVA database, which on average removed 90% of the reads, with the exception of samples C2 and IR2 for which 95% and 43% of the reads were retained, respectively, likely due to variation in the efficiency of experimental rRNA depletion [7]. Given the high number of reads retained after filtering for samples C2 and IR2 Table 1, a Cochran C test was performed to estimate significant differences in variance for any sample with respect to the entire group variance. The sample variance for C2 (0.9226) was significantly higher than the variance for the other samples (p-value of 2.2e-16) Table S2, potentially leading to biases. In consequence, the C2 sample was removed from subsequent analyses. For the IR2 sample, in which 43% of reads were retained, the calculated variance (0.6849) was not significantly different from the other samples.

**Table 1.** Reads counts (in millions) per sample through the preprocessing and mapping pipeline.

| Category | Features<sup>1</sup> | C1<sup>2</sup> | C2 | C3 | IR1 | IR2 | IR3 |
| --- | --- | --- | --- | --- | --- | --- | --- |
| Raw reads |  | 15.360 | 16.830 | 16.060 | 16.200 | 16.530 | 17.330 |
| After adapter removal |  | 15.360 | 16.820 | 16.050 | 16.200 | 16.520 | 17.320 |
| After rRNA removal |  | 1.820 | 16.090 | 2.290 | 1.300 | 6990 | 1.140 |
| Unmapped |  | 0.160 | 1.460 | 0.220 | 0.110 | 0.660 | 0.110 |
| Mapped | 3223 | 1.660 | 14.630 | 2.070 | 1.190 | 6.330 | 1.030 |
| *CDS*<sup>3</sup> | 3168 | 1.044 | 10.804 | 1.376 | 0.697 | 5.251 | 0.620 |
| *ncRNA*<sup>3</sup> | 1 | 0.079 | 0.539 | 0.090 | 0.049 | 0.299 | 0.034 |
| *rRNA*<sup>3</sup> | 7 | 0.036 | 0.005 | 0.045 | 0.059 | 0.034 | 0.073 |
| *tRNA*<sup>3</sup> | 47 | 0.004 | 0.123 | 0.005 | 0.005 | 0.010 | 0.003 |
| *Unassigned*<sup>3</sup> |  | 0.496 | 3.159 | 0.553 | 0.380 | 0.735 | 0.300 |

<sup>1</sup> Number of features identified in the annotated genome. <sup>2</sup> Values in Millions of reads. <sup>3</sup> Features belonging to mapped category.

Sequences were mapped against the reference *D. swuensis* genome DY59. The percentage of mapped reads ranged between 90.55% and 91.38%, with a maximum of 3 allowed mismatches Table 1. This range is expected when mapping against a different strain of the same species, due to intraspecific variability [10] [63] [57]. Features annotated as CDS, ncRNA, rRNA, and tRNAs were extracted from the dataset for differential expression analysis. The majority of the mapped reads corresponded to CDS (67.48 *±* 7.27%; mean *±* SD) distributed among 3168 genes. A total of 4.15 *±* 0.46% of the remaining reads mapped to a single ncRNA, making this the highest-scoring single feature. rRNA (7 features; 2.82 *±* 2.12%) and tRNAs (47 features; 0.37 *±* 0.15%) showed lower counts. Approximately 25.15 *±* 5.69% of mapped reads could not be assigned to any annotated feature Table 1.

As can be seen in Table 1, on average 10% of the reads failed to map against the reference genome. To identify the putative origin of these sequences, 10% of the unmapped reads (182,615 for controls and 86,952 for irradiated samples) were queried against the NCBI non-redundant (nr) nucleotide database using BLASTn. A total of 61,151 (33.49%) and 31,607 (36.35%) reads for controls and irradiated samples, respectively, were identified as having significant hits to the database (with an e-value threshold of 1e-5). Taxonomic assignment of the BLASTn results examined using MEGAN showed that for both controls and irradiated samples, *∼*31.15 *±* 0.19% of the reads corresponded to *Deinococcus*-related bacteria, another *∼*65.08 *±* 2.025% had no hit to the database, and the remaining 3.98 *±*0.18% were assigned to other bacterial groups Fig S1. The fact that *∼*30% out of the 10% unmapped reads were assigned to other *Deinococcus* species suggests intraspecific strain variation, in concordance, the alignment of these reads against reference genome DY59 (through Blast) recovered matches associated to described protein and RNA metabolism with identity values over 85.

### Differential expression analysis

To identify genes potentially involved in resistance to UV exposure, differential expression analyses were performed using all identified genomic features (CDS, ncRNA, rRNA, and tRNA) using control (C1 and C3) and irradiated samples (IR1, IR2 and IR3). Because an independence assumption is required to obtain accurate conclusions, it is essential to minimize external factors that could affect gene expression, regardless of the experimental condition being tested. The data were therefore first filtered by removing low count features (less than 5 CPM [counts per million]) and normalized by 1) sequencing depth and feature length variation (RPKM), and 2) taking into account sample total RNA content using the TMM method, (Trimmed Mean of M values is the average expression value after removing the most variant features of the data); this normalization takes into account sample-to-sample variation Fig 2A and Fig 2B.

**Fig 2.**
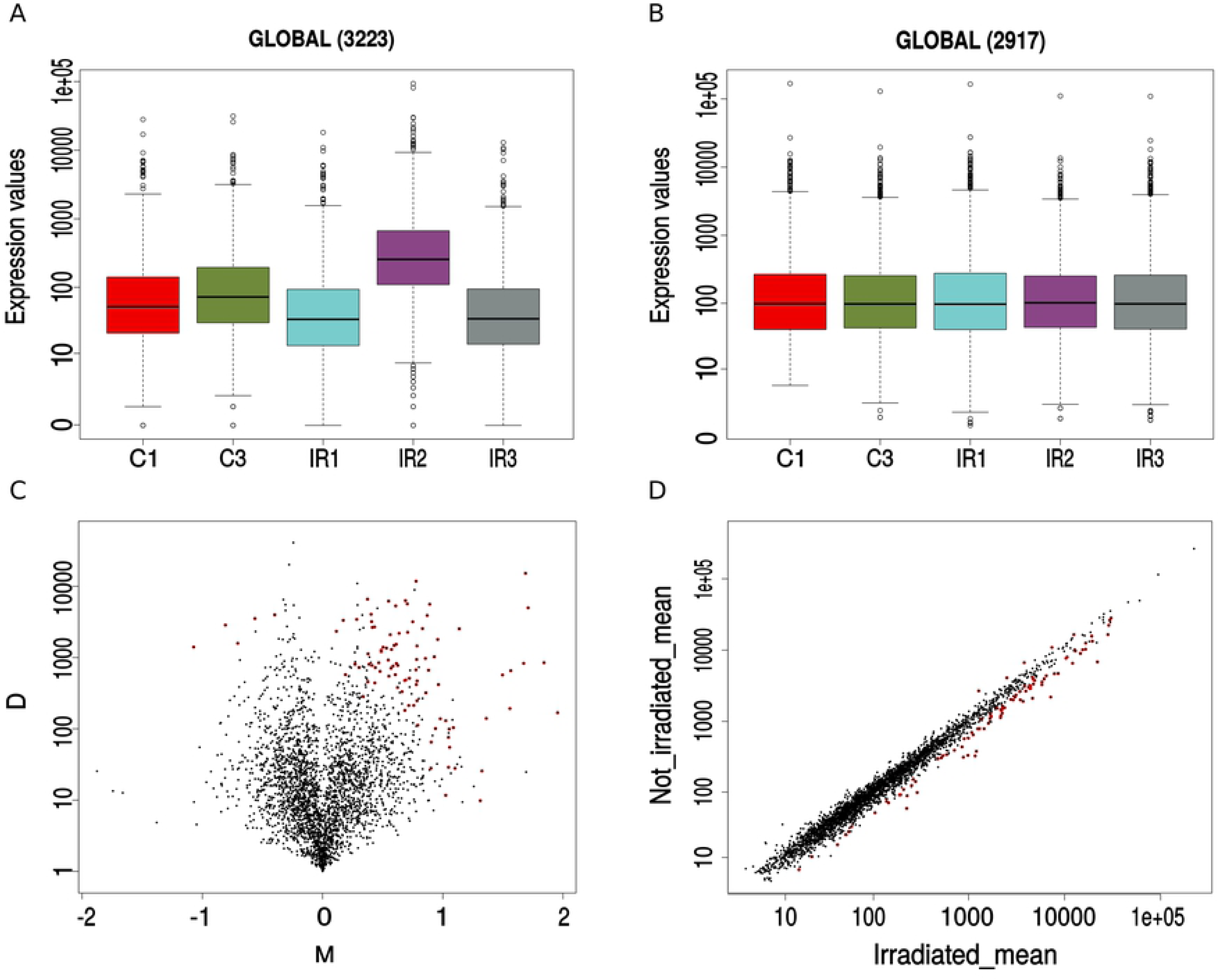
Differential expression analysis between irradiated and control samples. oxplots showing expression values (in counts per million) for control (C1, C3) and irradiated (IR1-IR3) samples before **(A)** and after **(B)** the filtering of low counts (CPM *<*5) and the normalization process done by RPKM (reads per kilobase of feature length per million mapped reads) and TMM (trimmed mean of M-Values). **(C)** Volcano plot of log-fold change (M) vs. the absolute value of the difference in expression between conditions; genes with a probability of differential expression *>*0.8 are shown in red, values of M *>*0 represent upregulated genes. **(D)** Correlation plot between irradiated (x-axis) and control (y-axis) mean expression. Genes deviating from expected with a probability *>*0.8 are shown in red. Values below and above the diagonal represent differentially expressed genes for the irradiated condition.

A total of 96 differentially expressed features with log_2_ fold-change values ranging between −1.07 and 1.95 and a probability for differential expression (p) *>*0.8 were obtained Fig 2C and Fig 2D. The chromosomal location of features that were up or down regulated did not show a particular position bias. The 96 detected features corresponded to four rRNAs, 13 tRNAs and 79 CDS (23 were hypothetical proteins), but only 14 CDS presented log_2_ fold-change values *>*1, indicating an over expression of at least twice as much as the control condition. Of these, 10 had functional annotation and 4 were hypothetical-proteins Table 2. The over-expressed CDS included genes for a GntR-like bacterial transcription factor, a proline dehydrogenase (key gene for homeostasis and ROS control in cells), RNA helicase (involved in ribosome biogenesis, initiation of translation), CrcB (protein for transmembrane transport of fluoride), an alpha/beta Hydrolase, a GTP-binding protein, Hemolysin, an ABC transporter ATP-binding protein and a pyr operon involved in synthesis of pyrimidines.

**Table 2.** Differentially expressed genes.

| GenID<sup>1</sup> | I_mean<sup>2</sup> | NI_mean<sup>2</sup> | Theta<sup>3</sup> | Prob<sup>4</sup> | Log<sub>2</sub>FC<sup>5</sup> | Function |
| --- | --- | --- | --- | --- | --- | --- |
| QR90_RS11755 | 224.78 | 57.99 | 0.98 | 0.92 | 1.95 | GntR |
| QR90_RS02510 | 984.31 | 332.59 | 1.24 | 0.9 | 1.57 | Hypothetical protein |
| QR90_RS11750 | 289.78 | 98.42 | 0.92 | 0.85 | 1.56 | Proline dehydrogenase |
| QR90_RS09640 | 226.93 | 88.31 | 1.11 | 0.92 | 1.36 | RNA helicase |
| QR90_RS15645 | 40.86 | 16.31 | 0.87 | 0.8 | 1.33 | CrcB protein |
| QR90_RS08275 | 14.76 | 5.95 | 0.65 | 0.81 | 1.31 | Alpha/beta hydrolase |
| QR90_RS05520 | 4636.4 | 2108.59 | 1.58 | 0.98 | 1.14 | 30S ribosomal protein S8 |
| QR90_RS09935 | 50.29 | 23.44 | 0.73 | 0.8 | 1.1 | Hypothetical protein |
| QR90_RS15125 | 194.2 | 91.37 | 0.82 | 0.84 | 1.09 | Hypothetical protein |
| QR90_RS06220 | 103.97 | 49.97 | 0.66 | 0.82 | 1.06 | GTP-binding protein |
| QR90_RS15620 | 54.06 | 26.16 | 0.78 | 0.83 | 1.05 | Hypothetical protein |
| QR90_RS03530 | 144.65 | 70 | 0.84 | 0.82 | 1.05 | Hemolysin |
| QR90_RS09365 | 21 | 10.33 | 0.68 | 0.82 | 1.02 | ABC ATP-binding protein |
| QR90_RS01280 | 253.16 | 124.44 | 0.74 | 0.8 | 1.02 | Bifunctional pyr operon |

<sup>1</sup> Only CDS with probability value over 0.8 and $\log_2$ fold-change $\geq 1$ are shown.
<sup>2</sup> Irradiated (I) and non-irradiated (NI) expression means.
<sup>3</sup> Theta: differential expression statistic calculated as $(M + D)/2$ , where M is the $\log_2$ -ratio of the two conditions and D is the difference in expression between conditions (including a correction for the biological variability of the corresponding feature).
<sup>4</sup> Prob: probability of mixed distribution (mixed because it is calculated from features changing between conditions and invariant features).
<sup>5</sup> $\log_2$ fold-change.

Given that some of the genes previously reported for *Deinococcus* strains as being involved in UVR resistance [63] [37], such as DNA repair mechanisms, pigment production and efflux pumps (for *Mn*^+2^ mainly), were not present among the most differentially expressed genes, a search for orthologues of radiation-resistance genes reported from *D. radiodurans* and *D. gobiensis* was performed. All twenty-seven reported genes (e.g., citB, ddrI, phoR, phrB and mutT) were recovered with significant e-values (*<*0.01) but with low identity values (between 30-50% at the DNA level) and a log_2_ fold-change value not significant between the conditions tested (maximum log_2_ fold-change 0.6). In consequence, those genes were not used for further analyses.

To further characterize the 23 differentially expressed hypothetical proteins, these were analyzed for possible functional domains through HMM (Hidden Markov Models) search against the pFam database. Several domains were identified, some of which were identified as related to photosystem II (PsbP), type III secretion system lipoprotein chaperone (YscW), copper chaperone pCu(A)C, WD domain (G-beta repeat), winged helix-turn helix, and DoxX categories. Four proteins were identified as containing conserved domains of unknown function (DUF) Table 3.

**Table 3.** Predicted domains present in hypothetical proteins reported by HMMER.

| GenID | Hits Found | Identifier<sup>1</sup> | Decription<sup>1</sup> | E-value<sup>1</sup> | Log<sub>2</sub>FC<sup>2</sup> |
| --- | --- | --- | --- | --- | --- |
| QR90_RS02510 | 1 | DUF4395 | Domain of unknown function | 5.2E-26 | 0.9 |
| QR90_RS15125 | 6 | WD40 | WD domain, G-beta repeat | 7.5E-06 | 0.84 |
| QR90_RS03320 | 1 | DUF456 | Domain of unknown function | 1.5E-35 | 0.81 |
| QR90_RS10725 | 1 | PsbP | PsbP | 7.6E-07 | 0.92 |
| QR90_RS08755 | 1 | HTH_33 | Winged helix-turn helix | 5E-14 | 0.92 |
| QR90_RS15830 | 1 | YscW | Type III secretion system | 1.6E-07 | 0.92 |
| QR90_RS03570 | 1 | DUF4384 | Domain of unknown function | 1.2E-10 | 0.91 |
| QR90_RS08995 | 1 | DUF1540 | Domain of Unknown Function | 8.2E-08 | 0.82 |
| QR90_RS02980 | 1 | PCuAC | Copper chaperone PCu(A)C | 8.4E-24 | 0.8 |
| QR90_RS11035 | 1 | DoxX | DoxX | 2.9E-10 | 0.87 |

<sup>1</sup> Only values for top-hit are shown.
<sup>2</sup> Log<sub>2</sub> fold-change.

### qRT-PCR validation

In order to validate the differential expression found with RNASeq, RT-qPCR was performed on three genes that had the highest log_2_ fold-change expression ratios: an RNA helicase (QR90 RS09640), a GntR family transcriptional regulator (QR90 RS11755) and the gene for proline dehydrogenase (QR90 RS11750). All three genes tested showed over 2-fold increase in expression (2.31, 2.14 and 2.05, respectively), thus confirming the observed RNA-seq data.

### Identification of ncRNAs

To identify additional differentially expressed features in the transcriptomes of UVC-exposed *D. swuensis* cells, and according to recently-proposed roles of ncRNAs in the rapid recovery after cellular stress, a de novo search for these regulatory entities was implemented [57]. Analysis of 3,355 intergenic regions from the D. swuensis reference genome retrieved 1,979 candidate ncRNA sequences. The criteria included a minimum cut-off for intergenic regions of 50bp and a minimum sequencing depth threshold of 6X, based on the mapping distribution, to eliminate regions with low coverage Fig S2. These candidates were compared against covariance models (CMs) built from the Rfam database. CMs are statistical models of structurally annotated RNA multiple sequence alignments that allow a flexible search for both primary and secondary RNA structures against a known dataset [4].

The CM search reported a total of 1,598 matches, but only 290 were below a search threshold of 0.1 (parameter that describes the number of hits one can “expect” to see by chance when searching a database). These significant matches were composed by 109 RNAs involved in post-transcriptional modification (such as snRNAs/snoRNAs), and 166 regulatory RNAs (including 97 miRNAs, 20 lncRNAs, 29 cis-regulatory elements, 15 antisense and 5 CRISPR RNAs). The remaining elements included one ribozyme, three antitoxin and 11 other RNA classes. Six candidates Table 4 were significantly related to small cytoplasmic Y RNAs (Rsm Y) (e-value *<*0.05). The log_2_ fold-change values for the differentially expressed ncRNAs oscillated between −1.03 (for mir-234) and 1.68 (for CsrC), which doesn’t indicate a tendency towards down or up-regulation under the UV-stress condition. However, the average of probability values for all ncRNAs was only 0.26*±* 0.23, whereas CsrC showed a probability of 0.79 Table 4. Although this probability is not equivalent to a p-value, the higher it is, the more likely that the difference in expression is due to the change in the experimental condition and not to chance.

**Table 4.** Significant ncRNA candidates reported by Infernal and log_2_ fold-change values for differential expression analysis.

| Category | Code | E-value | Model name<sup>4</sup> | GC<sup>5</sup> | Prob | Log<sub>2</sub>FC<sup>6</sup> |
| --- | --- | --- | --- | --- | --- | --- |
| Up<sup>1</sup> | nc0013 | 3.3E-05 | CsrC | 0.67 | 0.79 | 1.68 |
|  | nc0176 | 0.064 | Pxr | 0.5 | 0.45 | 1.46 |
|  | nc0001 | 0.074 | ar45 | 0.47 | 0.01 | 1.11 |
|  | nc0011 | 0.01 | mir-761 | 0.68 | 0.39 | 1.05 |
| Reported<sup>2</sup> | nc0989 | 5.9E-05 | RsmY | 0.7 | 0.20 | 0.77 |
|  | nc1573 | 0.00092 | RsmY | 0.68 | 0.03 | 0.59 |
|  | nc0919 | 0.052 | RsmX | 0.58 | 0.08 | 0.40 |
|  | nc0644 | 0.0059 | RsmY | 0.75 | 0.33 | 0.38 |
|  | nc0774 | 0.047 | RsmY | 0.67 | 0.04 | 0.27 |
|  | nc0445 | 0.018 | RsmY | 0.71 | 0.34 | 0.13 |
| Down<sup>3</sup> | nc0308 | 0.031 | mir-234 | 0.52 | 0.24 | -1.03 |

<sup>1</sup> Up: Upregulated ncRNAs, log<sub>2</sub> fold-change values are positive. <sup>2</sup> Reported: Significance and expression values for described RsmY family which were not differentially expressed in this study. <sup>3</sup> Down: Downregulated ncRNAs, log<sub>2</sub> fold-change values are negative. <sup>4</sup> ModelName: the name of the Rfam family/model of the hit. <sup>5</sup> GC: corresponds to the fraction of G+C of the hit. <sup>6</sup> Log<sub>2</sub> fold-change.

## Discussion

Natural ecosystems, and the organisms that inhabit them vary in their exposure to UVR. UVR determines the distribution and survival of microorganisms and consequently influences ecosystem dynamics and biogeochemical cycles [3]. From an evolutionary point of view, sensitivity to radiation indicates that UVR is an effective promoter of mutations, stimulating genomic variation, and could explain why high resistance is not widespread [20].

In this study, various isolates obtained from the *Espeletia* plant phyllosphere showed differences in resistance to UVR, indicating variability in their adaptability to UV exposure, despite being isolated from the same habitat. Experimentally observed differences in sensitivity to UVR have been reported for strains coming from different environments (related with their natural origin) [20], but similar levels of UVR resistance across different groups have been reported for a phyllosphere microbial community [54]. Our results indicate that habitat of origin is not necessarily indicative of resistance to extreme radiation, in this case UV-C, which seems to be affected by multiple factors in natural environments such as variable time and energy doses that are not necessarily reproduced in controlled laboratory experiments.

*D. swuensis* CG1225, with *>*60% survival at the highest dose tested (800*J/m*^2^), was the most resistant isolate recovered based on our radiation resistance experiments. The UV resistance of *Deinococcus spp.*, a group described as being “among the most radiation-resistant microorganisms that have been discovered”, has been widely recognized since 1956 [21]. Some *Deinococcus* members have shown tolerance to UV radiation (100 to 295 nm), and tolerance to ionizing radiation (5000 – 30000 Gy) [60]. As reference, 5Gy can destroy a human cell, 200-800 Gy will kill *Escherichia coli* and more than 4000 Gy breaks the resistance of tardigrades [60]. *Deinococcus* strains can resist ionizing radiation that is damaging to other organisms due to multiple mechanisms that can work synergistically to guarantee genomic integrity before the next cell division cycle [53]. Current reported strategies include efficient DNA repair (such as RecA, Pprl, Ppr), high antioxidant activities (CAT, SOD, POD, *Mn*^+2^), a unique cell structure (tetrad configuration for compartmentalization of DNA) [60], and protection of proteins [13]. Recent reports also indicate that ncRNAs may play a role in this resistance phenotype [57]. These organisms also have high genomic plasticity (i.e. high variability in genome content among closely related strains) and interspecies genomic diversity, when compared to the *Escherichia* genus, which can contribute to unique responses to environmental stress.

In this study, we used differential gene expression analysis to identify genes involved in the cellular response of *D. swuensis* CG1225 to UVR, a strategy which has been used to study other *Deinococcus* isolates [57] [44] [37] [63]. In contrast to previous studies in which treated cells were recovered after varying lengths of time, even up to three hours post treatment [63], here the *D. swuensis* CG1225 cells were harvested right after exposure and thus provide insight regarding the immediate response to UVR exposure and irradiation stress. This might explain why relatively few differentially expressed genes were identified, 14 CDS with log_2_ fold-change values *>*1 and a probability *>*0.8. Although functional domains with diverse biological functions were identified in these hypothetical proteins, none of them seemed to be associated with any known UV stress-responses Table 3. It is therefore unclear what the function of several of these proteins might be and how they may contribute to the UVR stress response. The predicted genes, however, were involved in global responses to stress, such as transcription regulation and transporters involved in cellular detoxification.

The overexpressed genes support the hypothesis of an organism that turns on its transcriptional machinery, in this case as an immediate response to prepare itself for the recovery of homeostasis as response to an environmental stressor. The highest log_2_ fold-change in expression was registered for a transcription factor belonging to the GntR family, which regulates several biological processes in diverse bacterial groups, however, details regarding its specific mechanism of action remain largely uncharacterized [15]. A few studies with mutants have shown that loss of function of this regulator protein is associated with a decrease in resistance to stress in *D. radiodurans* [15] and Bacillus subtilis [36]. However, the target genes for this transcriptional regulator remain elusive [1]. Other genes overexpressed under radiation exposure were an ABC transporter-system-related protein (ATP-binding protein), a *Mn*^+2^ transporter that has been shown to be key for ROS elimination [35] [63], and a hydrolase of the alpha/beta family. These last two genes can be potentially associated with cellular systems involved in cleaning toxic compounds produced during DNA repair. The activity of hydrolases modulating cellular redox processes has been described for many organisms [56] and in *D. radiodurans* it prevents incorporation of damaged nucleotides into DNA [1] [38]. These differentially expressed features indicate conditions that trigger the synthesis of genes and recycling of cellular components (such as chemical residues, oxidized nucleotides, etc.) from damaged biomolecules. Examples of such recycling mechanisms in *Deinococcus* come from studies showing that activity of Nudix-like hydrolases and RNA enzymes are essential for stress resistance [43] [1] [39] [38].

In this work we observed differences with respect to previous studies with *D. radiodurans* [37] and *D. gobiensis* [63]. In particular, we did not detect genes previously identified to be involved in resistance of *Deinococcus* strains, such as ddrA/ddrB genes for repair proteins, ddrC/ddrE/ddrP genes for damage response proteins and the fliY transporter. Neither these genes nor their orthologues were present among the differentially expressed genes. This discrepancy might indicate that different strategies may be involved regarding tolerance to radiation for *D. swuensis* CG1225 compared to both the widely studied *D. radiodurans* and *D. gobiensis* [42]. It could also reflect differences in experimental conditions, such as exposure to different levels of radiation [22] [14], the culture growth conditions and the amount of time allowed for cell recovery after UV exposure (from minutes to hours). In our work, the cells were exposed to a comparatively low level of radiation (considering the maximum level of resistance expressed by *D. swuensis*) and were harvested right after UVR treatment, thus providing an immediate snapshot, a “first quick response”, of the cellular response. In other studies, *D. radiodurans* transcriptome analysis has been conducted using longer recovery times, and even several hours, after exposure [63]. Finally, difference in results could also be due to genome variability. The high variability in genomic organization (genome size, number of chromosomes, plasmids, etc.) in *Deinococcus* sequenced isolates has been proposed as a potential source of interesting adaptations [21]. A comparison among *D. geothermalis, D gobiensis* and *D. proteolyticus* showed, for example, a core genome of 1369 genes and *∼*600-1700 accessory species-specific genes [21], which could harbor potential functional differences even among related species.

When analyzing the data obtained from the RNA-seq experiments, several reads could not be mapped to the reference genome used (*D. swuensis* DY59). The percentage of unmapped reads across the samples (*∼*10%) falls within the expected for RNA-seq experiments in which reads are mapped to a reference genome different from the evaluated isolate [63] [57], and is also consistent with the reported variability among *Deinococcus* genomes [21]. However, 30% of these unmapped reads showed identity values over 85% against the reference genome through a blast alignment, which reasserts the idea of intraspecific diversity for *Deinococcus sp*. Furthermore, an average of *∼*25% of the mapped reads failed to map within annotated features. Most of these fell near (50-100pb) to the start/end positions of annotated features, suggesting that they correspond to either transcribed but un-translated regions or miss-annotated features in the genome, a reasonable explanation due to the draft version of the available reference sequence.

Given the recent reports regarding the identification of differentially expressed ncRNAs in *Deinococcus* strains, we looked for these elements in our RNA-seq data. Even though the libraries were not experimentally enriched for short ncRNAs, an exploration of reads mapping to intergenic regions allowed the recovery of some well-represented families, which can be potentially involved in the irradiation response and have not been previously reported for the reference *D. swuensis* isolate [26]. The mechanism of action for ncRNAs in radiation response is a topic of current active research, and some recent studies suggest that ncRNAs, due to their small size, might remain largely undamaged by radiation and hence be the first responders, inducing and regulating cellular function recovery [57].

Several ncRNAs were identified in this study, some potentially involved in protecting against irradiation stress. These ncRNAs included members of the RsmY and RsmX families that bind and regulate molecules, such as translational proteins RsmA/CsrA and the sigma factor RpoS (a central regulator of the general stress response) [49] [23]. Previous experiments have shown that KsgA, which belongs to the RsmA family of ncRNAs, participates in the maintenance of translational fidelity under oxidative stress in *Staphylococcus aureus* [31]. CsrC, another promising ncRNA identified here, regulates the pleiotropic gene csrA (related to RsmY and RsmX) and can cause a decrease in oxidative stress resistance in Campylobacter jejuni when damaged [16]. Other ncRNAs identified corresponded to the Mir-761 Mir-234, ar45 and Pxr families, the reported functions for these ncRNA families do not have a clear relationship with radiation resistance; understanding their roles in resistance would require additional studies.

In summary, high-throughput sequencing of RNA provides a global view of the genomic responses and biological strategies required for cellular adaptation. Particularly, RNA-seq provides the possibility of measuring small-scale expression changes and can disclose novel cellular strategies responses [58]. Through RNA-seq, non-common overexpressed genes and novel ncRNAs families were reported for our *D. swuensis* strain isolated from the phyllosphere of plants. These findings offer an additional perspective for understanding bacterial resistance to radiation stress and expands our knowledge of bacterial transcriptomic dynamics.

## Conclusion

The transcriptional behavior of *D. swuensis* under the UVR stress condition studied here revealed differentially expressed genes that differ from mechanisms commonly reported for related species and expanded our understanding of UVR resistance in bacteria. The functions identified as relevant to the UVR response were involved in cell detoxification, regulation and reduction of stress by oxidation damage caused by ROS species. In addition, we also identified genes with undefined functions that may also play a role in radiation resistance. The ncRNA’s emerged as key players in the cellular response after UVR. Analysis of intergenic reads under covariance models retrieved families previously unannotated in *D. swuensis*, expanding its genomic annotation and uncovering ncRNA’s as key players in the cellular response after UVR. There was a tendency towards down-regulation of ncRNAs, an observation that needs further studies to validate and corroborate their role in the dynamic response after a radiation exposure event. This study contributes to the characterization of microbial biodiversity and provides evidence for new mechanisms involved in radiation-resistance. Describing and understanding these adaptations to extreme conditions may lead to potential applications, like preservation of products such as anticancer drugs and vaccines.

## Conflict of Interest Statement

The authors declare that the research was conducted in the absence of any commercial or financial relationships that could be construed as a potential conflict of interest.

## Author Contributions

Jorge Díaz-Rianõ designed and validated the computation pipeline, performed all the sequencing data processing including DE analysis, ncRNA search and primer design; led the data analyses and the manuscript preparation. Leonardo Posada provided input on bioinformatics pipeline and analysis and, with Catalina García-Castillo, performed *D. swuensis* survival curves, RNA extraction and RTqPCR validations. Carlos Ruiz and Ivan Acosta participated in the experimental design and sampling, performed the isolation of bacterial strains and the screen for UV resistance. Alejandro Reyes participated in the data analyses and manuscript preparation. Maria Mercedes Zambrano participated in the experimental design, provided support during the experimental and analytical stages, helped with data analysis and manuscript proofreading. No other conflict of interest relevant to this article was reported.

## Funding

This work was partially supported by Colciencias (Grant No. 657065843848) and was done under MADS contract no. 76-2013 for access to genetic resources.

## Acknowledgments

The authors thank Dra. Claudia Chica Pratesi for support in the analysis process and corrections about the analysis workflow. The authors also acknowledge the Universidad de Los Andes for providing a graduate assistance fellowship to JID and the High Performance Computing (HPC) Service that contributed to the research results reported within this work.

## Data Availability Statement

The datasets analyzed for this study can be found in the ENA (https://www.ebi.ac.uk/ena) under the accession number PRJEB33086.

## Supporting information

**S1 Fig.Taxonomic assignment of the BLASTn results for unmapped reads agaist nr database.** The reads were processed through MEGAN software and corresponds to controls and irradiated samples.

**S2 Fig.Histograms of read counts per sample mapping to intergenic regions.** Dotted line corresponds to the selected cutoff (log10 of 0.86) implying a minimum of 6 reads per region.

**S1 Table.Primers employed for qRT-PCR.** Three genes were used for evaluation, and one for normalization. T_*M*_ : Melting Temperature. GC%: Percent of G+C content

**S2 Table.Cochran’s C test for variance of groups:** C:ratio of the largest variance to the sum of the variances, n:number of observations (genes) in each group, k:number of groups.

